# Sequential screening nominates the Parkinson’s disease associated kinase LRRK2 as a regulator of Clathrin-mediated endocytosis

**DOI:** 10.1101/2020.04.28.053660

**Authors:** George R. Heaton, Natalie Landeck, Adamantios Mamais, Mike A Nalls, Jonathon Nixon-Abell, Ravindran Kumaran, Alexandra Beilina, Laura Pelligrini, Yan Li, International Parkinson Disease Genomics Consortium (IPDGC), Kirsten Harvey, Mark R. Cookson

## Abstract

Mutations in leucine-rich repeat kinase 2 (*LRRK2*) are an established cause of inherited Parkinson’s disease (PD). LRRK2 is expressed in both neurons and glia in the central nervous system, but its physiological function(s) in each of these cell types is uncertain. Through sequential screens, we report a functional interaction between LRRK2 and Clathrin adaptor protein complex 2 (AP2). Analysis of LRRK2 KO tissue revealed a significant dysregulation of AP2 complex components, suggesting LRRK2 may act upstream of AP2. In line with this hypothesis, expression of LRRK2 was found to modify recruitment and phosphorylation of AP2. Furthermore, expression of LRRK2 containing the R1441C pathogenic mutation resulted in impaired clathrin-mediated endocytosis (CME). A decrease in activity-dependent synaptic vesicle endocytosis was also observed in neurons harboring an endogenous R1441C LRRK2 mutation. Alongside *LRRK2*, several PD-associated genes intersect with membrane-trafficking pathways. To investigate the genetic association between Clathrin-trafficking and PD, we used polygenetic risk profiling from IPDGC genome wide association studies (GWAS) datasets. Clathrin-dependent endocytosis genes were found to be associated with PD across multiple cohorts, suggesting common variants at these loci represent a cumulative risk factor for disease. Taken together, these findings suggest CME is a LRRK2-mediated, PD relevant pathway.

## Introduction

PD is a common, age-dependent neurodegenerative disease characterized in part by nigral neuronal loss and motor symptoms. Mutations within the *LRRK2* gene have been shown to segregate with PD in large families.^1^ Additionally, common variants at the *LRRK2* locus have been linked to sporadic PD through GWAS.^2,3^ The *LRRK2* locus is therefore described a pleomorphic risk locus, containing both rare and common disease-linked variants.^4^ Importantly, PD arising from *LRRK2* mutations is indistinguishable from sporadic PD in terms of age of onset and clinical manifestation.^5,6^

The protein encoded by the *LRRK2* gene, Leucine-rich repeat kinase 2 (LRRK2), is a 286kDa multi domain protein belonging to the ROCO family.^7^ All ROCO proteins are characterised by a GTP binding domain (Ras of complex proteins, Roc), followed in tandem by a C-terminal of Roc domain (COR).^8^ Additionally, LRRK2 contains a kinase and multiple protein-protein interaction domains. These include amino-terminal armadillo and ankyrin repeats, followed by 13 leucine rich repeat regions (LRR) and 7 WD40 repeats at the C-terminus.^9,10^ To date, all mutations found to segregate with PD in families are clustered within the catalytic domains and have been found to alter the inherent biochemical properties of LRRK2. Three separate mutations have also been described at the 1441 position (R1441C/G/H) in the ROC domain, which have been shown to lower rates of GTP turnover.^11^ Because it is thought that the GTP bound form of LRRK2 is active in cells, diminished GTP hydrolysis would be expected to result in enhanced overall activity.^1,11^ Activity of the ROC-GTPase domain has also been shown to regulate kinase activity, dimerisation and LRRK2-dependent toxicity.^12^ The most common mutation, G2019S, is located within the kinase domain and directly leads to ~2 fold increase in kinase activity by increasing Vmax.^13^ Recent data suggests that small Rab GTPases may act downstream of LRRK2, with Rab29 also being proposed as an activator of LRRK2.^14–16^ Collectively, these results show that the various pathogenic mutations in LRRK2 result in gain of function that is associated with neuronal toxicity. Conversely, several studies have shown gene knockdown or pharmacological kinase inhibition can limit the detrimental effects of LRRK2 pathogenic mutations.^17–20^ These findings have prompted the development of LRRK2 targeting kinase inhibitors, which are currently in clinical trails. However, despite intensive study into the physiological function of LRRK2, a consensus on how LRRK2 acts within cells and how pathogenic mutations compromise this function has yet to be achieved.

Screening for binding partners has been a widely adopted strategy to narrow down the physiological activity of LRRK2.^21,22^ Given the genetic evidence implicating the ROC domain in PD pathogenesis, along with the general observation that GTPases often bind other proteins to either control their activity or to mediate cell signalling, we decided to focus on the mammalian LRRK2 ROC domain for identification of novel protein interactors of LRRK2. A subsequent siRNA screen was then used to determine whether any candidate interactors could modify subcellular LRRK2 localisation. We find that *AP2A1*, a gene coding for the α subunit of the adaptor protein 2 (AP2) complex, acts as a potent regulator of Rab29-dependent LRRK2 Golgi recruitment and kinase activation, which is a phenotype enhanced by pathogenic LRRK2 mutations.^21^ The AP2 complex is assembled into a heterotetrameric formation, containing two heavy subunits (α & β), one medium subunit (μ) and one small subunit (σ) and functions as a cytosolic bridge for cargo molecules and clathrin during the early stages of CME.^23^ Analysis of LRRK2 KO tissue further revealed a significant depletion of heavy and medial subunits of the AP2 complex alongside clathrin heavy chain. In cells, expression of the R1441C mutant was observed to induce dephosphorylation of AP2, reduced LRRK2-dependent AP2 clustering and slowed clathrin-dependent endocytosis. In neurons, LRRK2 R1441C knock in cells demonstrated a modest, but significantly reduced amount of synaptic vesicle endocytosis – suggesting mutation-dependent defects are conserved *in vivo*. Finally, using polygenic risk profiling we demonstrate an increased incidence of SNPs linked to Clathrin-dependent endocytosis genes in multiple PD patient cohorts, indicating this pathway is of direct relevance to disease aetiology.

## Results

### Overlapping screens nominate AP2 as a functional binding partner of LRRK2

We employed an unbiased protein-protein interaction screen to identify LRRK2 ROC domain binding partners (figure 1a & b). Recombinant wild-type and synthetic mutant ROC proteins were purified as described previously.^24^ GST-ROC (amino acids 1328 – 1516) constructs were transformed into modified *E. coli* and purified from the soluble fraction of bacterial lysates.^24^ All GST-fusion proteins were resolved and detected using SDS-PAGE and Coomassie stain, respectively (supplementary figure 1a). The GST-ROC protein was observed at an apparent molecular weight of ~46kDA. *In vitro*, the LRRK2-ROC domain selectively binds GTP and GDP nucleotides with similar affinities through a phosphate-binding P-loop motif.^12^ GTP binding was assayed to verify purified ROC retained its proposed biochemical activity. GSH-agarose beads with purified GST-fusion proteins were incubated with radiolabelled GTP and analysed for binding efficiency (supplementary figure 1b). Negligible GTP binding was detected for GST or for GST-ROC-K1347A, a synthetic ROC domain mutant with impaired GTP binding, compared to WT ROC.^25^ These results suggest that the isolated LRRK2 ROC domain is appropriately folded in solution and therefore a suitable probe for interactions screening.

**Figure 1.**
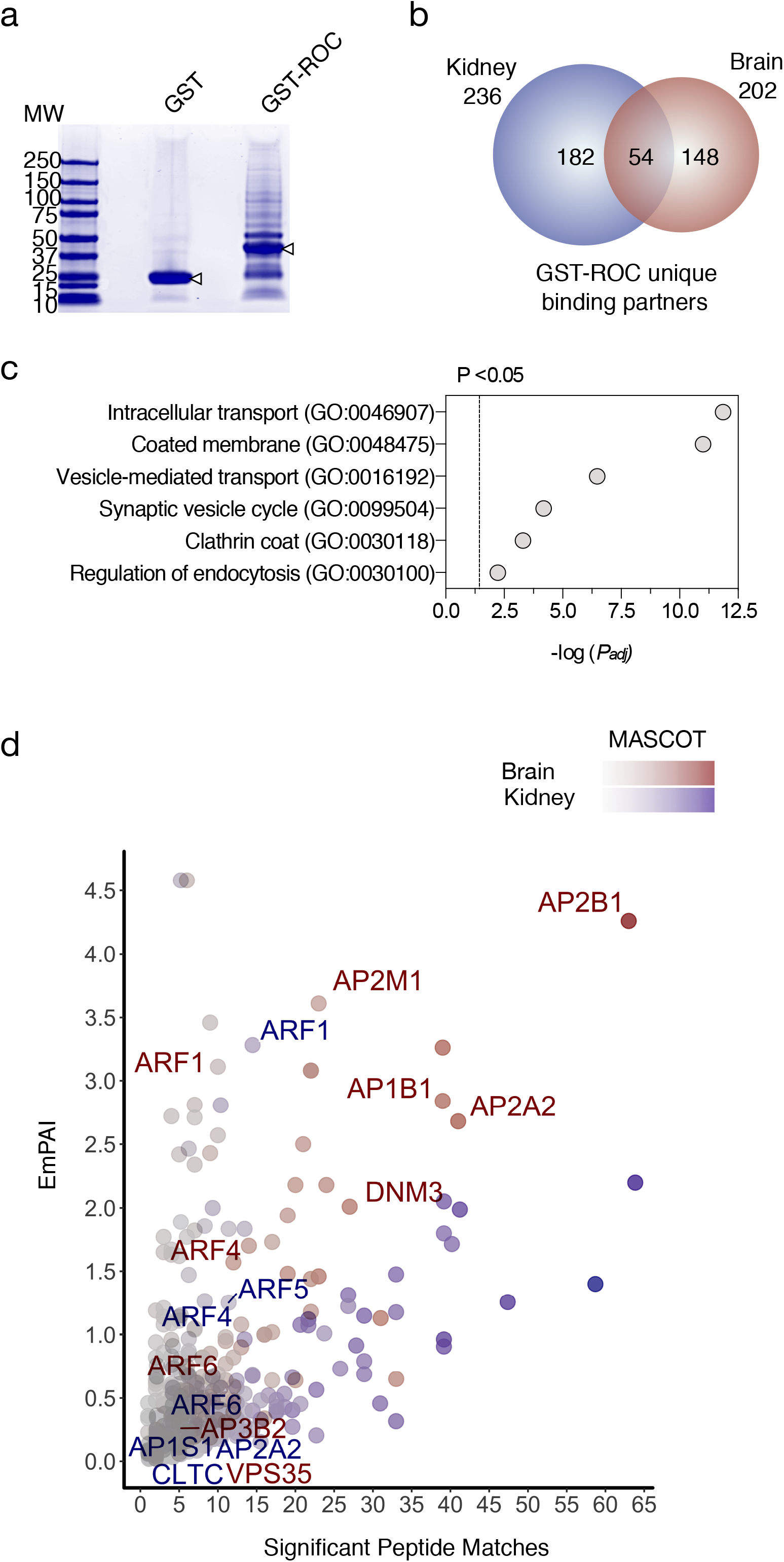
ROC domain protein-protein interaction screen. (a) Brain samples incubated with GST and GST-ROC resolved using SDS-PAGE and Coomassie stain to visualise protein bands. (b) Venn diagram summary of interacting proteins, colored by tissue used for analysis (blue – kidney, red – brain) and sized by number of proteins in each set. (c) Selected GO term enrichments from BP and CC categories using gProfileR. Results show - log P values below a threshold of p<0.05 after Bonferroni correction for multiple testing. (d) Scatter of massspectrometry results plotted by exponentially modified protein abundance index (EmPAI) and significant peptide matches. Blue points indicate kidney proteins, red represent brain and darker colors signify higher MASCOT scores. Annotated genes were found within GO nominated enrichments.

To identify LRRK2 binding partners, purified GST-ROC was incubated with soluble brain and kidney lysates from 1-year-old mice (figure 1). Approximately two hundred unique candidate ROC binding proteins were recovered from brain and two hundred and thirty six from kidney. Fifty-four proteins were shared between the two datasets (figure 1b). To better characterize these datasets, GO analysis was employed to identify discrete functional enrichments. A number of enrichments associated with intracellular transport mechanisms were nominated including the cellular component (CC) terms: ‘Clathrin coat’ (GO:0030118) and ‘Coated membrane’ (GO:0048475). Similarly, the Biological process (BP) terms: ‘Vesicle-mediated transport’ (GO:0016192), ‘Regulation of endocytosis’ (GO:0030100) and ‘Synaptic vesicle cycle’ (GO:0099504) were also deemed significant (Figure 1c). Several studies have demonstrated a link between LRRK2 activity and a variety of vesicular trafficking pathways making these a prime candidate for its cellular function.^21,26–30^ Interestingly, we identified several candidate interactors that were consistently nominated across these GO enrichments (figure 1d). These results suggest that vesicular trafficking proteins are authentic interactors of LRRK2.

To investigate whether any of the interacting partners or closely associated proteins recovered in our ROC screen could act as a modifier of LRRK2 in cells, we employed a targeted siRNA screen to examine localization of LRRK2 (figure 2 a&b). A functional interaction has been previously characterized between LRRK2 and the small GTPase RAB29.^16,31^ Transfection of RAB29 causes a robust relocalization of LRRK2 to the Trans-Golgi. Critically, this phenotype is exacerbated by pathogenic *LRRK2* mutation suggesting it may be of direct relevance to PD risk.^16^ We first generated an extended protein-protein interaction network to provide full coverage of clathrin-related proteins from publically available datasets (supplementary figure 2a) then generated a custom siRNA library against these targets. As positive and negative controls, siRNAs targeting CK1a and ARHGEF7 were included. For each condition, the percentage of cells with LRRK2-TGN localisation was normalised to nuclei counts. This value was expressed as a mean Z-score relative to WT LRRK2 treated with non-targeting control (NTC) siRNA. Mean-Z scores for each treatment condition were compared to NTC siRNA and corrected for multiple comparisons using bonferroni post-hoc test. In line with previous findings, depletion of CK1A increased LRRK2 localisation to the TGN, whereas knockdown of ARHGEF7 decreased the same parameter.^32^ Of 10 genes that had significant effects on LRRK2-mediated clustering, the α-subunit of clathrin adaptor protein complex 2 (*AP2A1*) had the largest effect (MeanZ, 4.1338, Adj P, 0.0002, figure 2c, supplementary figure 2b).

**Figure 2.**
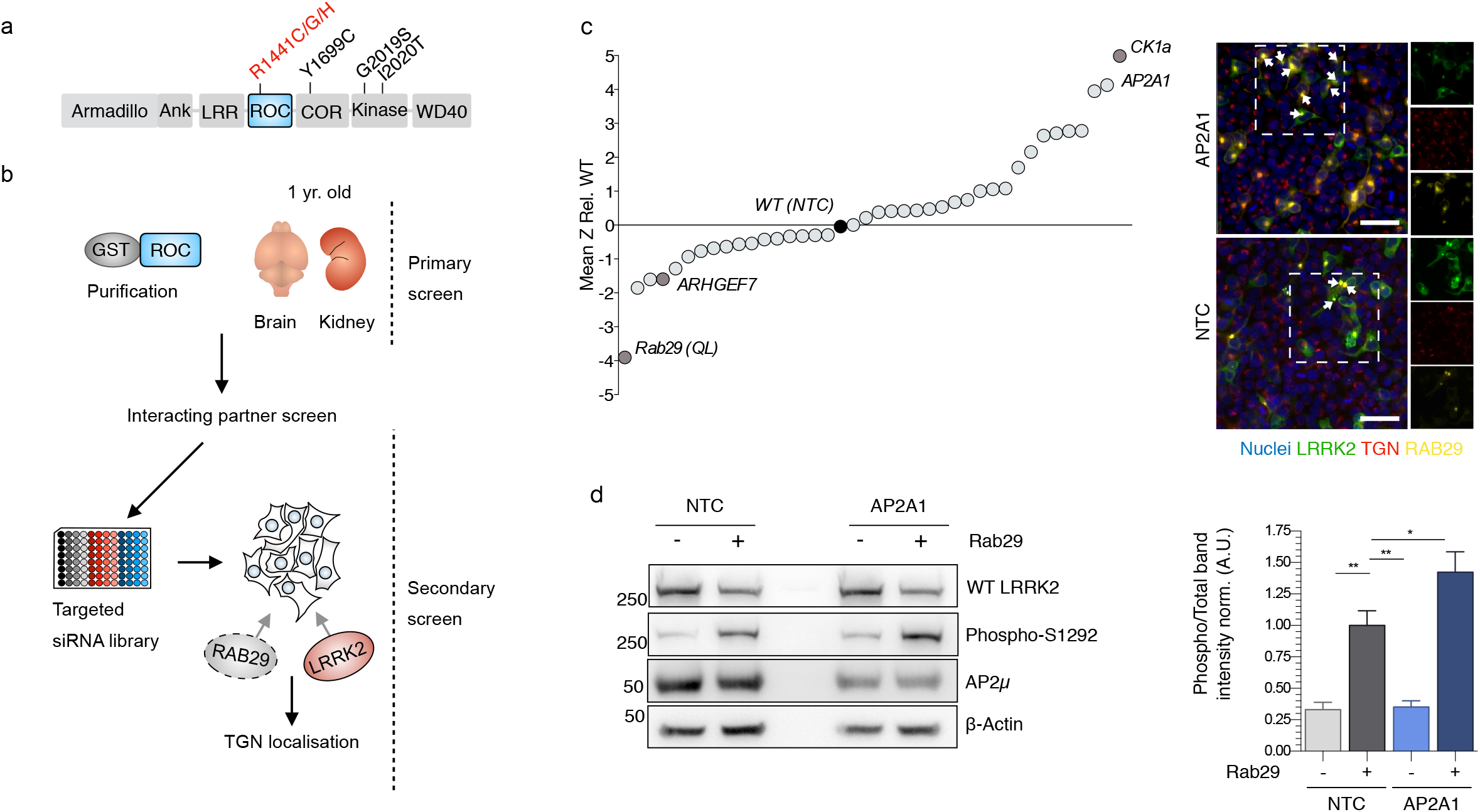
siRNA screen for modifiers LRRK2-TGN localization. (a) Diagrammatic representation of full length LRRK2 with PD associated mutations annotated. ROC domain is highlighted. (b) Outline of sequential screening strategy. Primary screen involved a protein-protein interaction screen using the purified ROC domain, a secondary screen assayed endogenous knockdown of selected candidates to assess modification of RAB29-dependent LRRK2 relocalisation. (c) Effects of knockdown of candidates on LRRK2 TGN localization. Mean Z scores from two independent screens were calculated relative to cells transfected with wild type RAB29 and treated with a non-targeting control siRNA (WT (NTC), black) and ranked according to effect size. Dark grey shaded points indicate positive controls. *AP2A1* emerged as a strong positive regulator of LRRK2 TGN relocalisation compare to non-tageting control siRNA (NTC) (MeanZ, 4.1338, AdjP, 0.0002. Scale bar indicates 50 μm and applies to all images). (d) AP2A and RAB29 both regulate LRRK2 S1292 autophosphorylation. Cells transfected with plasmid (-) or RAB29 (+) were also treated with siRNA against AP2A1 or a non-targeting control (NTC) were immunblotted for the indicated proteins. Markers on the left of all blots are in kilodaltons. *AP2A1* knock down following single transfection of LRRK2 did not significantly affect kinase activity. However, depletion of endogenous AP2A in the presence of RAB29 promotes kinase activity of LRRK2 (n = 6; AdjP, 0.0486, P < 0.05*; One-way ANOVA with Dunnett’s multiple comparisons test).

LRRK2-TGN localisation induced by RAB29 expression has been recently shown to stimulate kinase activity.^15^ Consistent with these observations, we observed co-expression of RAB29 and LRRK2 causes a significant increase in S1292 autophosphorylation (figure 2d). *AP2A1* knockdown following single transfection of LRRK2 did not affect kinase activity. However, depletion of endogenous *AP2A1* in the presence of RAB29 significantly increased kinase activity of LRRK2 (figure 2d). Therefore, AP2A1 knockdown both promotes LRRK2 relocalization and kinase activity in cells. For these reasons, a potential functional interaction between LRRK2 and the AP2 complex was explored in greater depth.

### LRRK2-AP2 association is conserved across tissue types and dysregulated in KO animals

As ranked by MASCOT score, AP2A2, AP2B1 and AP2M1 were all within the top 10% of LRRK2 interactors recovered from brain lysates. AP2A2 was present within the kidney mass spec results (supplementary table 3). In order to reliably detect the presence of AP2 subunits, we first validated antibodies using siRNA (supplementary figure 4). To confirm ROC-AP2 binding using an orthogonal technique, replicate pull downs of LRRK2 ROC were performed with kidney and brain lysates and samples were immunoblotted for specific AP2 subunits using these validated antibodies. Replicate pull-downs revealed discernable bands for each of the heavy and small subunits of AP2 in both tissues (figure 3a). Although the size of GST-ROC precluded detection of AP2μ using immunoblotting, this subunit was previously nominated as a ROC interactor by mass spec in brain (supplementary table 3).

**Figure 3.**
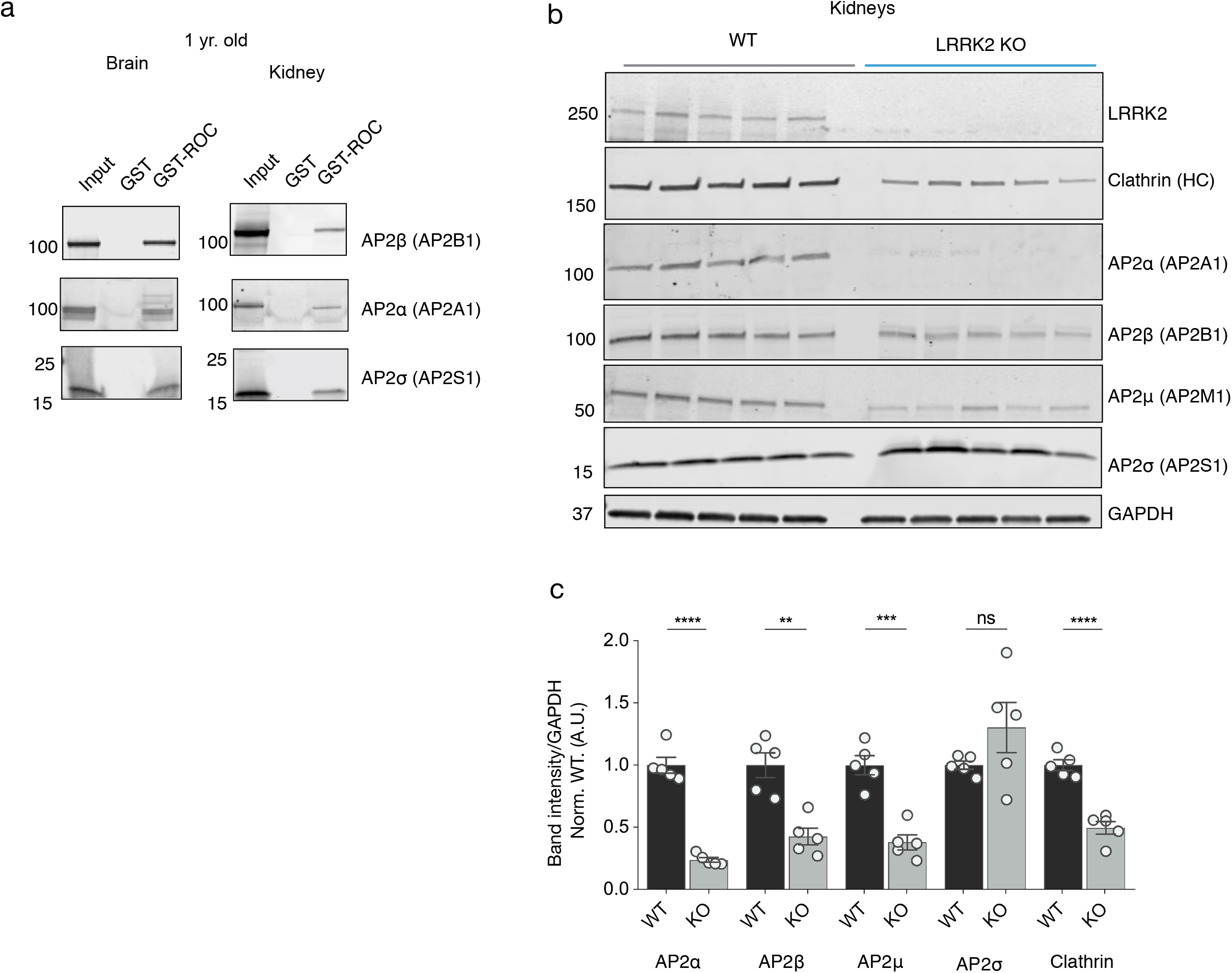
ROC binds to the AP2 complex in brain and kidney and is dysregulated in LRRK2 KO animals. (a) Validation of candidate hits by pull down assays with purified GST or GST-ROC protein from brain and kidney lysates. All organs were taken from 1 year old WT C57Bl6 mice. Western blots demonstrate binding of heavy and small subunits to the LRRK2 ROC domain in both tissues. Data is representative of *n* = 3 independent experiments with similar results performed in both tissues. (b) Deregulation of AP2 *in vivo*. Protein extracts from one yr. old WT or LRRK2 knockout kidneys were used for immunoblotting for the indicated proteins. Markers on the left of all blots are in kilodaltons. (c) Quantification of protein levels. All values are normalized to GAPDH as a loading control and expressed relative to the mean of wild type animals. (*n* = 5 animals; * P <0.05, ** P<0.01, Student’s t-test with Welch’s correction for unequal variance)

Though not previously detected by mass-spectrometry, immunoblots demonstrated robust binding of additional AP2 subunits to LRRK2 in tissue. Previous reported proteomic changes in LRRK2 KO kidneys prompted us to investigate levels of multiple AP2 components in these tissues.^33^ Kidney lysates from 1yr old. animals of WT and LRRK2 KO mice were processed as previously described in Pellegrini *et al*.^33^ Supernatant fractions were probed for all members of the AP2 complex as well as clathrin (figure 3b). Western blot analysis revealed a significant decrease in heavy and medium subunits of the AP2 complex (α, β & μ). Clathrin heavy chain was also significantly decreased in KO animals. Curiously, detected levels of the small subunit of AP2 (σ) were not significant between genotypes. These results confirm that LRRK2 physically interacts with the AP2 complex and that loss of LRRK2 results in lower levels of the AP2 complex.

### LRRK2 interacts with AP2 and mediates recruitment, phosphorylation and clathrin mediated endocytosis in a mutation-dependent manner

To investigate whether the observed regulation of LRRK2 activation is due to a physical interaction in cells, we purified the AP2 complex through expression and immunoprecipitation (IP) of AP2β. Western blot analysis detected all components of the AP2 complex as well as endogenous LRRK2, consistent with a strong physiological interaction (figure 4a). AP2 was also found to interact with co-expressed WT and the R1441C pathogenic variant of LRRK2 (supplementary figure 5a & b).

**Figure 4.**
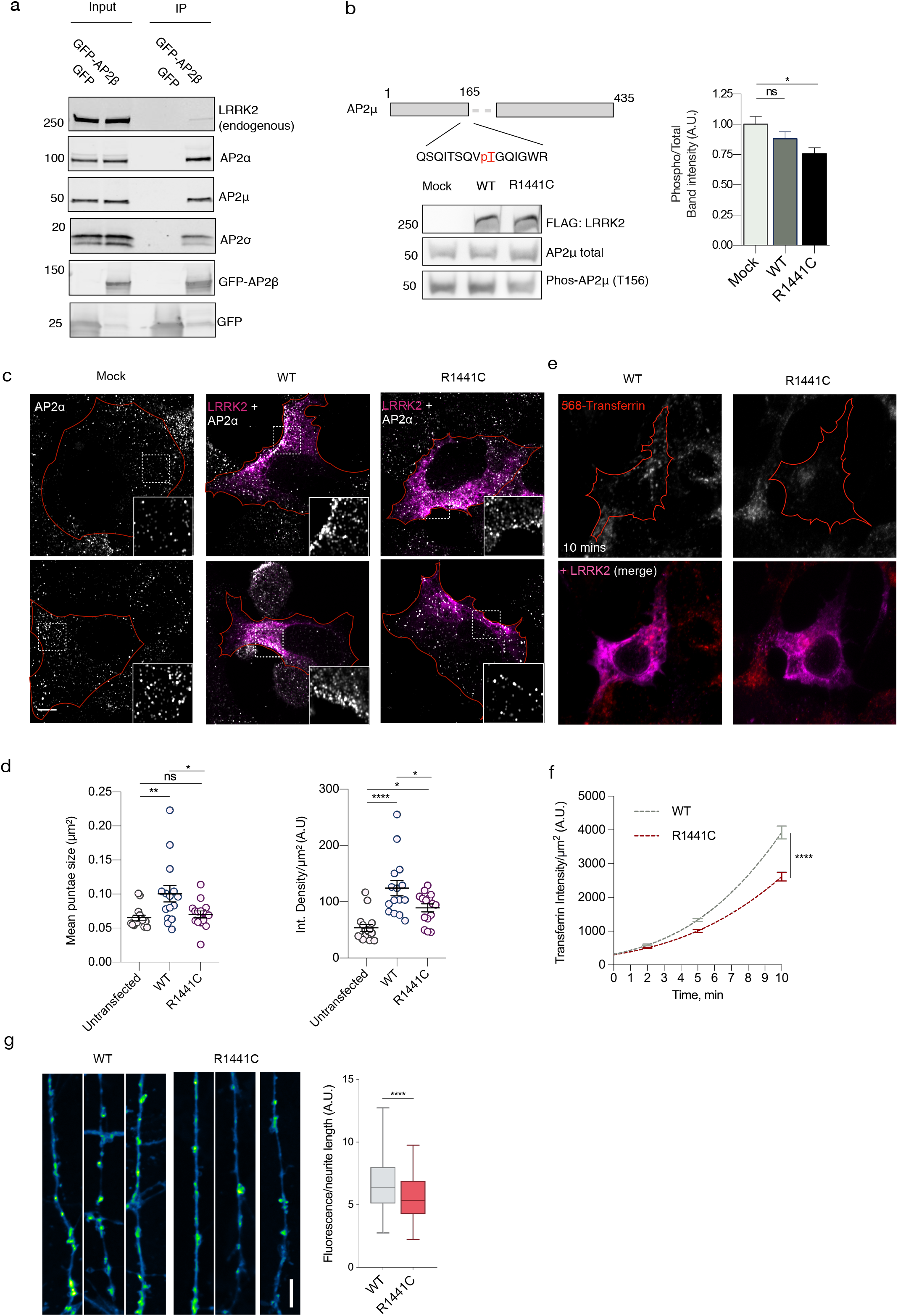
Cellular effects of LRRK2 on AP2. (a) We expressed and immunoprecipitated GFP or GFP-tagged AP2β subunit followed by immunoblot for endogenous members of the AP2 complex and for endogenous LRRK2. Markers on the left of all blots are in kilodaltons (b) R1441C LRRK2 lowers AP2μ phosphorylation. Mock transfected, or HEK293 cells overexpressing WT or R1441C LRRK2 were treated with pcalyculinA and probed for LRRK2, total and T156 phosphorylated AP2μ as indicated. Markers on the left of all blots are in kilodaltons. A significant decrease of phospho-AP2μ/total AP2 was observed in cells expressing R1441C LRRK2 (n = 9; AdjP, 0.0127, *P<0.05; One-way ANOVA with Dunnett’s multiple comparisons test). (c) Representative images of HEK293 cells stained for endogenous AP2α following either mock, LRRK2 WT or R1441C transfection. Scale bar: 5 μM. (d) Integrated density/area and mean vesicle size of AP2α was determined in areas positive for LRRK2 expression (ns, not significant; *, P<0.05; **, P<0.01; One-way ANOVA with Dunnett’s multiple comparisons test) (e) Representative images of HEK293 cells expressing either WT or R1441C LRRK2 and assayed for transferrin uptake at T10. (f) Time course quantification of internalised transferrin at 2,5 and 10 minutes. Expression of the R1441C mutant resulted in a significant decrease in internalised transferrin relative to WT (****, P<0.0001; Unpaired two-tailed t-test with Welch’s correction for unequal variance). (g) Representative images of FM 1-43 labelling in WT and R1441C primary hippocampal neurons cultured to DIV 14. Scale bar indicates 10 μm. Quantification of FM 1-43 intensity normalised to neurite length (****, P<0.0001; Unpaired two-tailed t-test with Welch’s correction for unequal variance)

The formation and internalisation of clathrin-coated vesicles during endocytosis is tightly regulated by protein kinases.^34^ More specifically, phosphorylation of AP2μ acts as a critical mediator of sorting motif binding and membrane association. ^35,36^ LRRK2 is a protein kinase, capable of phosphorylating itself and Rab proteins. Importantly, pathogenic mutations located within the GTPase domain have been shown to increase both LRRK2 autophosphorylation and phosphorylation of Rab substrates in cells. To independently verify this, we expressed increasing amount of wildtype LRRK2 and immunoblotted for phospho-Rab10 and LRRK2 S1292 autophosphorylation. As expected, S1292 levels relative to total LRRK2 remained constant but there was an increase in phospho-Rab 10 with higher LRRK2 levels (supplementary figure 5c). Expression of WT and R1441C LRRK2 at equivalent levels demonstrated mutation-dependent increase in phospho-rab10 (supplementary figure 5d). Having shown that expression of these LRRK2 constructs regulated known substrates as expected, we next asked whether AP2μ phosphorylation was influenced by LRRK2. The activity of the endogenous phosphatase PP2A makes phosphorylation of AP2μ transient. To assay phosphorylation of AP2μ, cells were treated with increasing concentrations of the PP2A inhibitor calyculin A, which increased phosphorylation of AP2μ in a dose-dependent manner (10-50nM, Ec50 = 19.95, supplementary figure 5e). Expression of R1441C LRRK2 lead to a decrease in AP2μ phosphorylation relative to WT or mock transfected cells in the presence of calyculin A (figure 4b). AP2μ is therefore unlikely to be a substrate for LRRK2 but LRRK2 mutations may indirectly affect AP2 function.

**Figure 5.**
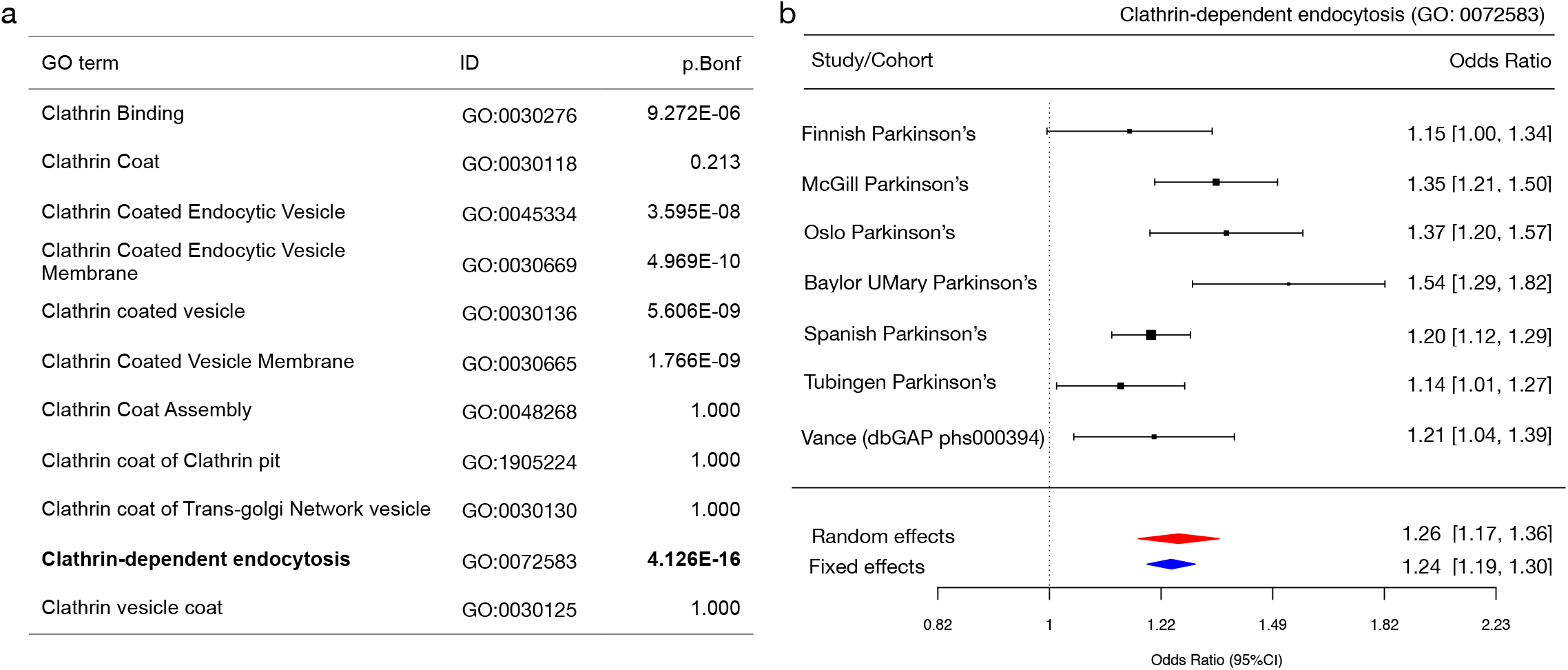
Polygenic risk scores (PRS) for clathrin-associated trafficking pathways. (a) Summary statistics for PRS estimates for each category and the number of SNPs analysed. P.Bonferonni denotes adjusted p values. (b) Forest plot of PRS scores for ‘GO: 0072583, Clathrin-dependent endocytosis. The size of each square at the point estimate of Odds Ratio is proportional to the number of samples used in each study. 95% confidence intervals are represented as horizonal error bars. Random and fixed effects meta-analysis of odds ratios are shown as red and blue diamonds respectively.

To further investigate the cellular effects of LRRK2 expression on AP2, we analyzed the subcellular AP2 distribution using fluorescent microscopy. The specificity of staining for endogenous AP2α was validated using siRNA treated cells (supplementary figure 6). In their nascent state, AP2 vesicles are broadly distributed throughout the cell, fluctuating between membrane-bound and a cytosolic pool during cycles of endocytosis. Once vesicles are uncoated, AP2 is recycled back to the cytosol. High-resolution Airyscan imaging of HEK293 cells revealed a broad distribution of AP2 throughout the cytosol with little distinct membrane localization (figure 4c). Following expression of WT LRRK2, we observed highly localized clustering of AP2 towards the outer cell edges and plasma membrane. Areas of positive LRRK2 expression showed significantly more clustering of AP2 punctae as reflected by increased fluorescence intensity and mean vesicle size. Remarkably, LRRK2-dependent clustering of AP2 was significantly lower with expression of the R1441C mutant (figure 4c & d). No significant differences were recorded in vesicle size between R1441C and mock-transfected cells. Some increase in staining intensity was observed between R1441C and untransfected cells, though both conditions were significantly lower than in cells transfected with WT LRRK2.

Recruitment of the AP2 complex to the plasma membrane occurs via binding of *μ* and α subunits to the membrane lipid PIP2 (phosphatidylinositol (4,5)-bisphosphate). PIP2-AP2 interactions facilitate the recruitment of clathrin triskellions to initiate the formation of a coated pit. ^23^ Endocytic cargoes containing tyrosine-based sorting motifs, such as the transferrin receptor (TfR), activate the AP2 complex to promote clathrin-coat assembly.^23^ Therefore, the dynamics of AP2 recruitment is essential in the initiation of CME. To understand the functional consequence of LRRK2 expression on CME, we first used a well-established transferin uptake assay. Transferrin is an iron-binding glycoprotein that enters the cell by clathrin-mediated internalisation of the TfR. For this reason, assaying uptake of fluorescently labelled transferrin molecules acts as a physiological readout for CME. Cells transfected with WT or R1441C LRRK2 were incubated with human transferrin labelled with Alexa-568 (Tf-568). Quantification of internalised cellular Tf-568 showed that R1441C LRRK2 significantly slowed the rate of transferrin internalisation, with the largest difference observed at the 10-minute time point (figure 5e & f). Taken together, these findings suggest the R1441C pathogenic mutation inhibits LRRK2-dependent recruitment of AP2 and impairs CME. To understand the role of endogenous wildtype LRRK2 in CME, the Tf-568 internalisation was assayed in LRRK2 depleted cells. NTC were compared to *AP2* and *LRRK2* siRNA treated cells. In line with reported data, loss of AP2 caused a significant decrease in internalised transferrin.^38^ Importantly, this result was mirrored by depletion of endogenous LRRK2 (supplementary figure 7).

These results suggest that the R1441C mutant impairs CME, via an AP2-dependent mechanism in transfected cell lines. To determine whether these effects are conserved in a more physiological model system, we investigated the impact of the same LRRK2 mutation in cultured hippocampal neurons from R1441C knockin mice compared to wild type controls. To establish that all signaling components were present in these preparations, we stained for endogenous AP2. Fluorescent microscopy revealed punctate staining throughout the neuronal cell body and neurites (supplementary figure 8a). Expression of LRRK2 in primary neurons similarly demonstrated a broad cytoplasmic distribution across all neuronal compartments (supplementary figure 8a). Primary neurons were also positive for the synaptic markers synaptophysin and PSD-95, indicating functional maturity by DIV 14. Immunoblot analysis also identified that these cells express endogenous LRRK2 and AP2 complex components (supplementary figure 8b). Taken together, these results support a potential physiological role for LRRK2 and AP2 within mature neurons.

A substantial body of evidence has suggested CME plays a fundamental role in the retrieval of vesicles within the synaptic terminal.^39,40^ In order to study CME in the context of synaptic recycling, we employed a well-established FM 1-43 stryl dye endocytosis assay.^41^ Primary hippocampal neurons derived from WT and R1441C knockin mice were cultured to DIV 14. Neurons were stimulated and incubated with FM1-43FX dye to label endocytic vesicles. Quantification of FM 1-43 fluorescence intensity per length of neurite indicated that endocytosis was moderately decreased in R1441C neurons (figure 4g). Critically, our findings indicate mutation-dependent defects in CME are conserved at endogenously levels of LRRK2 protein expression.

### Polygenic risk profiling nominates Clathrin associated pathways as a risk factor for Parkinson’s disease

Abundant literature suggest that *LRRK2*, as well as several other PD-associated genes, are key mediators of endocytosis and intracellular trafficking.^27,29,30,42–45^ However, it remains unclear if these pathways are causal for PD in humans. To investigate whether genes associated with clathrin-trafficking pathways contribute to heritable risk of PD, polygenic risk scores were calculated from GWAS datasets using GO terms for clathrin and related pathways.^46^ Of the 11 GO terms analysed, 6 were found to be significant after correction for multiple comparisons. Clathrin-dependent endocytosis (GO: 0072583) proved to be the most significant (P value = 4.126E-16, SE = 0.031) and had an odds ratio of 1.25 (figure 5a & b). The directionality of effect was consistent between different population datasets (figure 5b supplementary & table 9).

## Discussion

Although mutations in *LRRK2* are well established as causal for PD, the mechanisms by which these mutations contribute to cellular dysfunction and disease are incompletely understood.^47^ LRRK2 has been previously suggested to regulate many facets of intracellular trafficking including, synaptic vesicle endocytosis and recycling, trafficking and degradation of EGFR, actin remodeling on endosomes, trafficking of mannose-6-phosphoate receptors from the TGN to lysosomes and the recycling of receptors from the membrane to the TGN through the retromer complex.^26,27,30,33,42,48–50^ However, the mechanistic details underpinning these observations remain uncertain with competing substrates and protein interactors nominated to mediate cellular effects.

Here, through a sequential screening approach, we identify a functional interaction between LRRK2 and the AP2 complex that we confirmed in both kidney and brain. Although LRRK2 deficiency does not result in brain defects, LRRK2 KO kidneys exhibit progressive changes in autophagic activity accompanied by increased apoptosis and age-dependent renal pathology.^23,51^ Recent proteomic analysis of aged LRRK2 KO kidneys characterised a number of changes to lysosomal, cytoskeletal and protein-translation related proteins.^33,52^ Relative to their WT counterparts, aged KO animals also demonstrated a significant reduction in clathrin and AP2 heavy and medial subunits, α, β and μ respectively. These findings indicate AP2 is downregulated or degraded in the absence of LRRK2, although the precise mechanism by which this occurs is uncertain at this time. While endocytosis and autophagy are often described as separate entities, there is significant cross talk between these cellular pathways.^53^ Given this, it is plausible that endocytosis defects arising from the absence of AP2 components could contribute to the pathological changes observed in kidneys.

Interestingly, the pathological kidney phenotype first described in LRRK2 KO mice, has also been observed in RAB29 KO animals suggesting these two proteins work in a single pathway.^54^ In cells, RAB29 has been shown to bind LRRK2 to increase TGN-localisation and kinase activity.^15,21,55^ Our siRNA screen nominated *AP2A1* as a strong modifier of LRRK2 recruitment and showed that depletion of *AP2A1* in the presence of RAB29 caused a further increase of LRRK2 autophosphorylation. The mechanism by which this occurs is not yet known. One possible explanation is that RAB29 competes with existing interacting partners of LRRK2. Therefore, knockdown of LRRK2 binding partners may increase the cytosolic pool of LRRK2 available for RAB29 mediated activation and sequestration at the Golgi. Interestingly, knockdown of *AP2A1* was observed to cause reduced levels of the AP2μ subunit and may reflect AP2 complex instability. Recruitment of AP2 to the membrane is mediated by binding of AP2α to PIP2 and AP2μ interactions with cellular cargo.^56^ It is therefore possible that the loss of AP2 membrane associations is the principle factor causing increased translocation of LRRK2 to the TGN.

The primary function of AP2 is to bind to cargo molecules and recruit clathrin during the initial stages of endocytosis. Activity of the AP2 complex is tightly regulated through phosphorylation including at threonine (Thr) 156 of AP2μ.^35,36^ Critically, phosphorylated AP2μ exhibits a much greater affinity to sorting motifs relative to dephosphorylated forms.^36^ Inhibiting of AP2μ phosphorylation *in vitro* has been shown to prevent internalisation of transferrin into coated pits.^35^ Similarly, expression of phospho-null AP2μ has been shown to inhibit transferrin uptake in cells.^57^ These observations have lead to the hypothesis that continuous endocytosis requires the dynamic cycling of AP2 through phosphorylation and dephosphorylation – facilitating sequential rounds of cargo binding and internalisation.

We examined how LRRK2 expression affects AP2μ phosphorylation and found that expression of the R1441C mutant led to a significant decrease in AP2μ phosphorylation levels. This is in contrast to the increase in Rab10 phosphorylation that is thought to reflect the inherent gain of function caused by this mutation. It is feasible that these two observations may be mechanistically related. Rab5, a reported LRRK2 substrate has been proposed to play a role in uncoating of AP2 and clathrin from coated vesicles.^58,59^ Recruitment of the Rab5 guanine nucleotide exchange factor hRME-6 to CCV has been shown to cause dephosphorylation of AP2μ through displacement of the AP2-kinase AAK1.^58^ We speculate that phosphorylated Rab5 and perhaps other Rabs may be unable to cycle between endosomal compartments after Lrrk2-dependent phosphorylation, leading to downstream alterations in phosphorylation of proteins at clathrin coated vesiclesIf correct, this posits that while LRRK2 mutations are gain of function in terms of immediate biochemical effects, they may also have some features that appear to be a loss of function.

Such conjecture would be consistent with the observations that both overexpression of R1441C LRRK2 or depletion of endogenous LRRK2 significantly lowered the rate of transferrin internalization in cells as a measure of CME. We also demonstrate that neurons harboring a pathogenic LRRK2 mutation at the endogenous level demonstrated a significantly reduced amount of synaptic vesicle retrieval. This observation further supports prior studies that have demonstrated a reduction of CME in iPSC-derived LRRK2 R1441C and G2019S dopaminergic neurons as well as a significant decrease in several CME-associated proteins, including AP2.^28,60^ Collectively, our data is most compatible with several previous studies suggesting that either too much or too little LRRK2 can deregulate cellular activity.^26–29,48^

To date, GWAS have identified up to 90 independent PD-associated signals across 78 loci.^46^ Each of these signals carries a relatively small effect size and collectively account for 26-36% of PD heritability.^46^ As such, a significant proportion of PD-heritability is still unaccounted for, likely due to small effect sizes of undiscovered loci that do not surpass GWAS significance thresholds.^61^ Here, we used polygenic risk scores to investigate the potential contribution of clathrin-trafficking genes to PD risk. Four genes were present in all PD-associated categories: *AP2A1, AP2A2, SNAP91* and *PICALM* and two of these genes encode for isoforms of the α subunit of the AP2 complex. This analysis indicates that genetic variability within a subset of clathrin-associated genes is linked with an increased risk of PD and reinforce the concept that AP2 dysfunction may play a causal role in human PD.

## Conclusions

In summary, this study explores the role of LRRK2 in vesicular trafficking through identification of protein-protein interactions. We suggest AP2-LRRK2 interactions are important for the regulation of clathrin-mediated endocytosis and that LRRK2 pathogenic mutation can functionally impair this pathway in cellular models. These findings contribute to the current understanding of LRRK2 biology and suggest dysfunction of clathin-mediated endocytosis is relevant to PD pathogenesis.

## Methods and Materials

### GST-ROC purification and pull down assay

pGEX_GST, GST-ROC (1328-1516) and GST-ROC K1347N mutant in were each expressed in BL21 *E. Coli* and grown on antibiotic supplemented LB-Agar plates. Transformed cultures were grown at 37°C with constant shaking in 200mL of LB bacterial growth medium (KD medical) supplemented with antibiotic. When OD_600_ reached ~0.5, protein expressed was induced with Isopropyl β-D-1-thiogalactopyranoside (IPTG) added to a 200μM concentration and incubated for a further 2hrs. Bacterial cells were harvested by centrifugation at 1,500g for 10 minutes. Bacteria pellet was resuspended in 10mL of ice cold lysis buffer (50mM Tris-HCL, 50mM NaCl, 0.5mM Ethylenediaminetetraacetic acid (EDTA), 5% glycerol, 1mM Dithiothreitol (DTT), 0.1% Triton X-100, 5mM MgCl_2_., 5mM ATP and protease inhibitors cocktail). Lysis was aided using two cycles of pulse sonication (Fisher Scientific) for 10 minutes on ice. Cellular debris was pelleted by centrifugation at 12,000 rpm for 15 minutes. Resultant supernatant was incubated with 100μL of Glutathione Sepharose beads (GE healthcare) overnight with constant mixing at 4°C. Beads were spun down at 100g for 1 minute and washed 4x in ice cold lysis buffer and 1x in PBS and stored at 4°C.

Frozen adult (1 year) mouse Brains and Kidneys were homogenised using a 1mL Tissue Grinder (Wheaton) in 20mM HEPES tissue lysis buffer with protease inhibitors and centrifuged at 1,000g for 10 minutes. Triton X-100 was added to a 1% final concentration and samples were incubated on ice for 15 minutes before a final centrifugation at 21,000 rpm for 30 minutes in a tabletop centrifuge. Protein levels were normalised across samples using a Pierce™ BCA protein assay kit (ThermoFisher) according to manufacturers instructions. Tissue lysis supernatant was incubated with GST fusion proteins pre-coupled to Glutathione Sepharose beads for 30 minutes at room temperature. Beads were spun down at 1,000rpm for 1 minute and washed 3x in tissue lysis buffer with 1% Titron X-100. Samples were processed for western blotting using SDS-PAGE sample buffer (BioRad) and boiled for 5 minutes. For western blot analysis Kidney samples were prepared as described previously.^33^ In brief, kidneys were dissected from 1-year-old mice and frozen prior to preparation and homogenized in HEPES lysis buffer as described above. Homogenates were centrifuged at 300g for 3 minutes, supernatant was collected and centrifuged at 12,000g for 8 minutes – all centrifugation steps were performed at 4°C. Supernatant was collected and processed for Westernblot analysis.

### GTP binding

equal amount of GST, GST-ROC WT and GST-ROC K1347A mutant protein bound to Glutathione Sepharose 4B beads (GE Healthcare) were washed twice with Buffer A (20 mm Tris–HCl pH 7.5, 100 mm NaCl, 5 mm MgCl_2_, 1 mm NaH_2_PO_4_, 2 mm DTT). Beads were incubated overnight on ice in Buffer A containing 0.1 *μ*m ^3^H-GTP. Beads were then washed twice in Buffer A to remove unbound nucleotides, added to the Bio-safe II (RPI) scintillation cocktail and binding quantified using scintillation counting for H3 on an Tri-Carb 2810TR scintillation counter (Perkin Elmer).

### *SDS PAGE* and Western Blotting

Proteins samples were run on 4–20% Criterion TGX pre-cast gels (Biorad) in SDS/Tris-glycine running buffer and transferred to PVDF membranes by semi-dry trans-Blot Turbo system (Biorad). Membranes were blocked with either Odyssey Blocking Buffer (Licor Cat #927-40000) or 5% skimmed milk in TBST buffer (SIGMA) and incubated for 1h at room temperature or overnight at 4°. The membranes were washed with three 5 minute wash cycles in TBST at room temperature (RT) followed by incubation for 1h at RT with goat anti-mouse or rabbit IR Dye 680 or 800 antibodies (LICOR) or HRP-conjugated anti-mouse or rabbit secondary antibodies (Cambridge bioscience). Blots were washed as above at RT and scanned on an ODYSSEY^®^ CLx (Licor). For HRP-secondaries, each membrane is soaked in SuperSignal^®^ West Fempto/Pico maximum sensitivity substrate (equal parts Luminol and stable peroxide buffer, Thermo Scientific). Membranes were imaged using a GeneGenome XRQ Chemilluminescence imager (high quantium efficiency (QE) camera – Syngene). Quantitation of western blots was performed using Image Studio (Licor) or GeneSnap software.

### Cell culture

HEK293FT cells (Thermo Scientific) were maintained in Dulbecco modified eagle media (DMEM) containing 4.5 g L^−1^ glucose, 2 mM l-glutamine, and supplemented with 10% fetal bovine serum and 1% Pen Strep antibiotic (Thermo). Cells were maintained at 37°C in 5% CO_2_. All procedures with animals followed the guidelines approved by the Institutional Animal Care and Use Committee of National Institute on Aging. Primary neuronal cultures were prepared from newborn P0 C57Bl/6J mice. Dissected hippocampi were incubated in 10mL Basal medium eagle (BME) (Sigma) supplemented with 5ml papain solution (Worthington) for 30 minutes at 37°C. Brains were then incubated with 5μg of DNAseI and titurated to dissociate single cells. Cells were washed with two cycles of 10mL BME and counted. Cells were plated at ~ 0.5×10^6^ on Poly-D-lysine & laminine precoated coverslips in BME supplemented with B27, N2, 1mM Glutamax (Invitrogen), 0.45% glucose (SIGMA). Media was replaced the next day with BME supplemented with 2.5μM cytosine arabinose to kill off glial cells.

### Transfections

HEK293FT cells were transfected using Lipofectamine 2000 (Thermo Scientific), according to the manufacturer’s instructions. For siRNAs, cells were transfected with the SMARTpool or Individual ON-TARGETplus or NONtargeting scrambled siRNA using DharmaFECT I transfection reagent (Dhamacon Horizon discovery) according to the manufacturer’s instructions.

### LRRK2-RAB29 activation screen

Screen candidates were ordered on a custom pooled ON-TARGETplus siRNA library (Dharmacon). HEK293FT cells were seeded at a density of 100,000 per well within a 96 well plate and transfected with siRNA at a final concentration of 50nM. CK1A, ARHGEF7 and non-targeting control siRNA were seeded on each plate as positive and negative controls respectively. After 24 hrs, cells were transfected with 3xFLAG WT LRRK2 and 2xmyc WT Rab29 or 2xmyc Q67L Rab29 using Lipofectamine 2000 as per manufacturers instructions (Thermo Scientific). 30 hours following DNA transfection, cells were fixed with 4% paraformaldehyde (PFA) in PBS for 15 minutes at room temperature. Cells were left in PBS at 4°C overnight. Blocking and permeabilisation was performed with 5% FBS in PBS containing 0.1% Triton, all antibody incubation steps were performed in this solution. Cells were incubated with primary antibodies, M2-FLAG (Sigma) at 1:500, Myc (Chromotek) at 1:500, TGN46 at 1:1000 (Bio-Rad), for two hours at RT. Cells were washed 3x in PBS before incubation with secondary antibodies, Goat anti-Mouse Alexa Fluor 488, Goat anti-Rat Alexa Fluor 568, and Donkey anti-Sheep Alexa Fluor 647, all at 1:500 along with Hoechst 33342 (Thermo Scientific) at 1:10,000. Plates were imaged using a Cellomics VTI Arrayscanner at 20x objective with percentage of transfected cells showing positive colocalisation of FLAG WT LRRK2 and TGN46. Initial Pooled siRNA screen was run in duplicate, as was individual siRNA as part of POOL deconvolution. All values were normalized to cell number and compared to non-targeting siRNA control.

### Immunoprecipitation

HEK293FT cells were transfected using Lipofectamine 2000 (thermo) or FuGENE HD Transfection reagent (promega) according to manufactures instructions. Cells were lysed in IP buffer (20 mM Tris-HCl pH 7.5, 150 mM NaCl, 1 mM EDTA, 0.3% Triton X-100, 10% Glycerol, 1x Halt phosphatase inhibitor cocktail (Thermo Scientific)) and protease inhibitor cocktail (Roche) for 30 min on ice with constant rotation. Cell lysates were cleared at 20,000g for 10 minutes at 4°C in a tabletop centrifuge. Lysate supernatant was incubated with GFP-Trap agarose beads (ChromoTek) for 1 hour at 4°C with constant rotation. Beads were washed in IP wash buffer (20 mM Tris-HCl pH 7.5, 150 mM NaCl, 1 mM EDTA, 0.1% Triton X-100, 10% Glycerol) for four cycles. Following washing, sample volume was made up to 50μl and processed for Western blotting.

### Tissue processing

WT and KO kidneys were processed as previously described by Pellegrini *et al*.^33^ Frozen kidneys were homogenised using a gl in tissue lysis buffer (0.225M mannitol, 0.05M sucrose, 0.0005M HEPES, 1mM EDTA). Kidney homogenated was centrifuged in a tabletop centrifuge (Eppendorf) at 2000g for 3 minutes at 4°C. Supernatant was transferred to a separate microcentrifuge tube and pellet resuspended in tissue lysis buffer. Supernatent from the resuspended pellet was combined with the previous supernatant and centrifuged to clear lysates at 12,000g for 8 minutes. Supernatent was transferred to a new micro centrifuge tube and pellet discarded. One final ultracentrifugation step was performed on the supernatant at 55,000g for 30 minutes at 4°C. Supernatent was collected and processed for SDS-PAGE and western blot.

### Immunostaining

HEK293FT cells or primary cultures were fixed and blocked as described above. Primary antibodies were incubated for overnight at 4°C, washed three times in PBS for 5 minutes each and incubated with secondary antibodies as described above. Cover slips were mounted using ProLong^®^ Gold antifade reagent (Thermo Scientific).

### FM1-43 dye uptake assay

Primary neurons were cultured until day 14 in supplemented BME as described above. FM1-43FX stryl dye (ThermoScientific) was solubulised in HBSS at 37°C to a stock concentration of 1mg mL^−1^. FM1-43 dye uptake was performed as described previously.^41^ In brief, cells were washed briefly and then incubated at 37°C with HBSS media containing Ca2+ for 5 minutes. Media was aspirated and replaced with depolarising solution: HBSS containing 60mM KCL+ along with 5μg mL^−1^ FM 1-43FX stryl dye for 1 minute. Depolarising solution was replaced with HBSS with Ca2+ containing 5ug mL^−1^ of FM 1-43FX stryl dye for an additional 10 minutes during the recovery phase. Following incubation, cells were washed with constant perfusion with 3 mL with HBSS without Ca2+ in three cycles. Cells were fixed using 4% PFA and mounted prior to imaging as described above. A minimum of 6 mice per genotype were used to culture hippocampal neurons. Neurons were seeded at 0.5×10^6^ cells per well across 5 coverslips per genotype per experiment. Two independent litters were used for these experiments.

### Transferrin-568 internalisation assay

HEK cells were seeded at 0.3×10^6^ within a twenty-four well plate and either reverse transfected with siRNA or transfected after 24 hours with DNA constructs (as described above). 48 hours post seeding, Complete DMEM media was aspirated and cells were washed briefly in PBS and incubated with prewarmed serum-free media for 45 minutes at 37°C. This stage ensures cellular transferrin stores are depleted. Human transferrin-568 (ThermoFisher) was added at 10μg mL^−1^ concentration in serum free media and incubated at 37°C to allow for internalization. After sufficient time had elapsed, transferrin containing media was aspirated from cells, wells were washed twice in ice-cold HBSS to halt internalisation and fixed in 4% PFA for 15 minutes at room temperature. Cells were processed for immunocytochemistry and imaged as described above. When possible, plates were covered with foil post fixation.

### Microscopy

Confocal microscopy was performed using a Zeiss LSM 710 or Zeiss LSM 880. Higher resolution imaging was performed using a Zeiss 880 fitted with an Airyscan module. Data was collected using a 63×1.4NM objective under oil immersion (Carl Zeiss). Airyscan processing was performed using the Airyscan module in ZEN software package (Carl Zeiss).

### Image analysis

Quantification of images was performed using Fiji software (NIH). To determine AP2α integrated density per μm^2^, images were opened as split channels. LRRK2 expression channel was converted to 8-bit and a binary mask was created to outline the area of positive staining. The binary mask from the LRRK2 channel was imposed on the AP2α channel and measured for integrated density and area. To calculate particle size all surrounding area outside of the selection was cleared from the image and converted to 8-bit. The image was thresholded to a set value and converted to a binary mask. Finally, the image was inverted and analysed for particles to calculate the mean vesicle size per cell. For mock-transfected cells an identical workflow was used except ROI was manually approximated. To calculate transferrin internalisation, cells positively expressing LRRK2 were determined by eye within the 488-nm (green) channel. Brightness and contrast were adjusted so the whole cell could be visualized. An ROI was manually drawn around all edges of the cell. Integrated density was measured within the 568-nm (red) channel and expressed as function of the selection area. To quantify transferrin uptake in siRNA treated cells, images were opened as split channels, nuclei staining was converted into binary and selected to highlight individual cells. The selection area was expanded by 3 spatial units to cover perinuclear area and a portion of the cytoplasm. The selection area was imposed on the transferrin uptake channel and measured for integrated density and area. As above, these values were divided by one another and plotted per image. For FM1-43 dye uptake assay in primary neurons, all images were blinded and manually sorted to determine which were appropriate for quantification. This was determined by overall culture health how discernable individual neurites were. To quantify fluorescence, individual processes were selected using ‘Simple Neurite Tracer’ plugin in Fiji. Individual neurites were traced and selections exported as an ROI. Each selection was expanded by 3 pixel units and measured for integrated density and expressed as a function of its length.

### IPDGC data and Polygenetic risk score analysis

Quality control procedure within IPDGC GWAS datasets was as previously described.^46^ Briefly, samples with a call rate of <95% and whose X chromosome heterogeneity did not match clinical data were not included. Individuals with heterozygosity greater than six standard deviations from the population mean and those of non-European ancestry were excluded. All individuals related at more than the level of first cousin were excluded.

For PRS calculations, the same workflow as in the most recent PD GWAS-meta analysis,^46^ with the exception that only clathrin-associated GO terms were selected. PRS analysis was conducted using the R package PRSice2.^62^ Linkage disequilibrium (LD) pruning is a process of filtering SNPs so only those that are representative of genetic haplotype blocks are retained. LD clumping was applied using default settings (window size = 250 kb; r^2^ > 0.1). Empirical P values are generated through permutation testing whereby cases and control (segregated by cohort) are swapped in the withheld sample. 10,000 permutations were used to generate empirical P values ranging from 5×10^−8^ to 1×10^−4^ with increments of 5×10^−8^. Summary statistics were meta-analysed using random and fixed effects per study-specific dataset. For more details of included cohorts, see supplementary table 9.

### Bioinformatics and Statistical analysis

In all experiments, n represents the number of measures used in quantification. In mass-spec analysis was performed using the Mascot Server (version 2.5) to identify matching peptides in the Sprot Mouse Database (Uniprot Proteome ID: UP000000589). R was used to analyse results from LRRK2-TGN siRNA screen. Statistical analysis for experiments with only two treatment groups used Student’s t-tests with Welch’s correction for unequal variance. Outliers were identified with ROUT with Q set to 1%. For multiple groups, statistical significance was determined using one-way ANOVA with Bonferroni post-hoc test for multiple group comparisons. Dunnett’s multiple comparison test was used in instances where all groups were compared with a single control group. Comparisons were considered statistically significant where *p*<0.05. Data was plotted using Prism 7 (Graphpad) or R studio (https://www.rstudio.com/) and displayed as a mean with standard error of the mean (SEM) error bars.

### Antibodies and chemicals

The following antibodies used for immunoblot as described above are as follows: mouse monoclonal antibody to FLAG peptide (Sigma #F1804, 1:5000), Rabbit monoclonal to c-Myc (abcam #ab32072, 1:2000), Rabbit polyclonal to Alpha Adaptin (Proteintech #11401-1-AP, 1:2000), Mouse monoclonal to Alpha Adaptin (Thermo #MA1-064, 1:1000), Rabbit monoclonal to AP2M1 (Abcam #ab75995, 1:2000), Rabbit monoclonal to AP2S1 (Abcam #ab128950, 1:2000), Rabbit monoclonal to AP2B1 (Abcam #ab129168, 1:1000), R Mouse monoclonal to Anti- β-actin (Sigma #A5316, 1:3000), Rabbit polyclonal to GAPDH (abcam #ab9485, 1:2000), Rabbit monoclonal to LRRK2 (Abcam #MJFF2, ab133474, 1:1000), Mouse monoclonal to GFP (Roche #11814460001, 1:2000), Rabbit monoclonal to LRRK2 phospho-S1292 (Abcam #ab203181, 1:1000), Rabbit monoclonal to total-Rab10 (Abcam #ab237703), Rabbit monoclonal to Rab10 phospho-T73 (Abcam #ab230261, 1:2000), Rabbit monoclonal to AP2M1 phospho-T156 (Abcam #ab109397, 1:2000). The following antibodies used for immunocytochemistry as described above are as follows: mouse monoclonal antibody to FLAG peptide (Sigma #F1804, 1:500), sheep polyclonal to TGN46 (Biorad #AHP500G, 1:1000), rat monoclonal to myc-tag (Chromotek #9e1, 1:500), mouse monoclonal to Alpha Adaptin (Thermo #MA1-064, 1:500), mouse monoclonal to Transferrin Receptor (Thermo #H68.4, 1:500). Calyculin A protein phosphatase inhibitor (ab141784) was dissolved in DMSO and used at concentrations ranging from 10 to 50nM.

## Acknowledgments

We thank David Gershlick for kindly providing the GFP-AP2β plasmid. We would also like to express our thanks to Zhenyu Yue for allowing us to complete this study in his laboratory. We would like to thank all of the subjects who donated their time and biological samples to be part of this study. We would also like to thank all members of the International Parkinson Disease Genomics Consortium (IPDGC). For a complete overview of members, acknowledgements and funding, please see the Supplemental data and/or http://pdgenetics.org/partners.

## Author contributions

GH, KH and MC conceived and designed the majority of experiments. GH performed majority of experiments. GH, KH and MC analyzed majority of the data and wrote the manuscript. RK, JNA, NL and AM performed imaging experiments. MN, performed genetic analysis and analysed data. LP helped with tissue lysis experiments. AB performed GTP binding experiments. YL performed mass spectrometry. All authors read and approved the final version.

## Availability of data and materials

All raw data are available on request.

