## Supplementary data for "Sequential screening nominates the Parkinson’s disease associated kinase LRRK2 as a regulator of Clathrin-mediated endocytosis"

**Supplementary Figure 1. GST purification and GTP binding** (a) Representative Coomassie stained gels of purifications. Equal amounts of GSH-beads were resolved. From left to right, (b) GST-WT ROC domain showed significantly more GTP binding capacity relative to the GST-K1347A, an artificial mutant with lower GTP binding capacity, and GST as measured by counts per minute (CPMB) (one-way ANOVA with Dunnett's multiple comparison test, n = 3 technical replicates, **** P<0.001).

**Supplementary figure 2. Network and summary statistics** (a) Informatic extension to initial nominated protein candidates. Proteins associated with vesicular trafficking pathways were queried against the Intact database. Blue colours indicate those which were initially nominated as LRRK2 ROC domain interactors. Outlined are proteins known to form a single complex. (b) siRNA screen summary statistics. Values for individual screen Z-scores, mean Z and adjusted P value. Significance was determined by analysis of noramlised MeanZ scores and calculated using student's t-test with bonferroni post-hoc correction for multiple testing. Dotted line indicates significance threshold. Significance was determined using R. (c) Uncropped representative images of cells treated with NTC and AP2A1 targeting siRNA as part of a TGN-LRRK2 localisation screen (scale bar indicates 50 *μ*M and applies to all images).

**Supplementary table 3. Summary of mass spec results candidate protein interactors.** Proteins recovered in ROC interactor screen from brain and kidney datasets as ranked by MASCOT score.

**Supplementary figure 4. AP2 antibody validation.** HEK293 cells were treated with single siRNAs targeting genes of interest, numbers in brackets correspond to the molecular weight. All samples showed a significant drop in signal. (n = 4, Student's t-test with Welch's correction, P<0.01 ** P <0.05 *)

**Supplementary figure 5. LRRK2 is a cellular kinase that interacts with AP2. CalyculinA treatment increases AP2µ phosphorylation in a dose-dependent manner**. (a) Interaction of GFP-AP2β or GFP with co-expression of Flag-tagged LRRK2 (b) GFP IP of GFP-AP2β with co-expression of Flag-tagged WT or R1441C LRRK2. (c) Western blot analysis of increasing amounts of WT LRRK2 expression in HEK293 cells demonstrates an increase in phospho-Rab10/total (d) Expression of WT and R1441C LRRK2 in equal amounts causes a mutation-dependent increase in phospho-rab10/total rab 10 (n = 3, student's t-test with Welch's correction, ** P < 0.01). (e) Dose-dependent increase in AP2µ phosphorylation following calyculin A treatment of HEK293 cells (n = 3). (f) Representative images of HEK293 cells stained for endogenous AP2α following LRRK2 WT or R1441C transfection. Scale bar: 5 *μ*m. (g) Quantificaiton of the total levels of AP2 α fluorescence in LRRK2 transfected cells and normalised to cell area (student's t-test with Welch's correction, ns P < 0.4). (h) Representative images of HEK293 cells stained for LRRK2 and AP2S1 following overexpression of both proteins. Scale bar: 5 *μ*m.

**Supplementary figure 6. Immunocytochemistry validation of AP2 staining.** HEK293 cells were treated with NTC and *AP2A1* targeting siRNA. Depletion of AP2A1 caused a reduction of endogenous AP2α puntae. Scale bar: 10$\mu$m.

**Supplementary figure 7. Loss of AP2M1 and LRRK2 inhibits transferrin uptake** (a) Western blot analysis of siRNA treated HEK cells. Quantification demonstrates robust knockdown of target genes. All values are normalised to β-actin as a loading control. (b) Representative images of siRNA treated cells analysed for transferrin uptake at T=10. Unbiased automated quantification of transferrin fluorescence intensity/perinuclear area (n = 15 images, across 3 independent repeats. T = 10min, One-way ANOVA, Dunnett's multiple comparisons test, *** P <0.001,).

**Supplementary figure 8** (a) Representative images of primary hippocampal neurons expressing WT LRRK2 or stained for endogenous AP2α, synaptophysin, PSD95 or β-III Tubulin at DIV 14. (b) Immunoblot of primary hippocampal neurons at DIV 14.

**Supplementary table 9. Clinical and demographic characteristics of IPDGC data.** The IPDGC GWAS data set is comprised of 9983 individuals (5516 cases and 4467 controls) of European ancestry (see below).

Supplementary Figure 1.

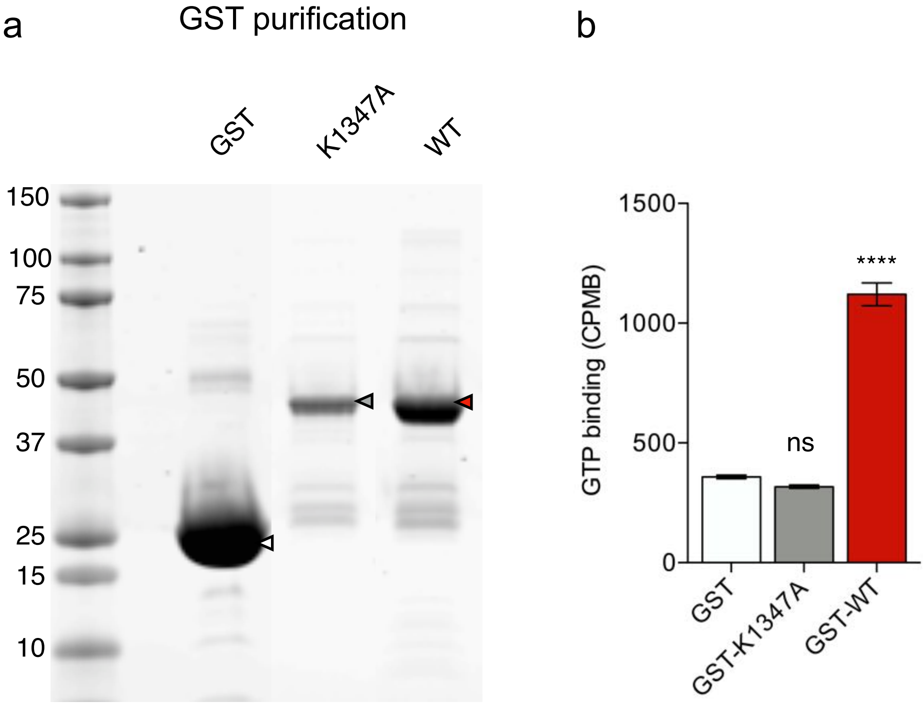

Supplementary Figure 2.

a

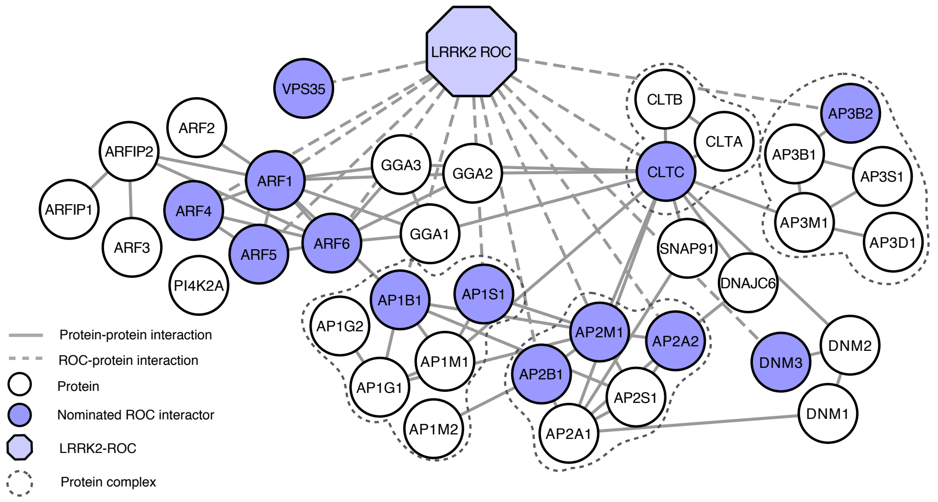


b

| Gene | MeanZ | Screen 1 Z | Screen 2 Z | Adj. P |
|---|---|---|---|---|
| QL | -3.9069 | -2.8979 | -4.8974 | 2.8000E-54 |
| CK1a | 4.9899 | 2.7705 | 7.1299 | 9.2200E-37 |
| ARHGEF7 | -1.5981 | -1.0429 | -2.1533 | 1.4800E-17 |
| AP1B1 | 2.7821 | 2.1685 | 3.3190 | 8.4200E-06 |
| ARF4 | 2.7109 | 2.4184 | 3.0034 | 9.5900E-06 |
| ARF6 | 2.6403 | 2.4203 | 2.8602 | 2.4800E-05 |
| AP2A1 | 4.1338 | 2.2270 | 6.0405 | 0.0002 |
| ARF1 | -1.8558 | -1.0660 | -2.6457 | 0.0006 |
| AP2A2 | 1.6937 | 1.7247 | 1.6666 | 0.0009 |
| DNM2 | 2.7760 | 1.4091 | 4.1430 | 0.0010 |
| AP1M1 | 3.9483 | 1.5947 | 6.3019 | 0.0031 |
| ARF3 | -1.6050 | -0.9495 | -2.1786 | 0.0058 |
| AP2M1 | 2.1563 | 0.7550 | 3.5577 | 0.0305 |
| GGA2 | 1.0825 | 0.6445 | 1.5205 | 0.1469 |
| AP3S1 | -0.7647 | -0.7529 | -0.7782 | 0.2751 |
| AP1M2 | 1.0052 | 0.6268 | 1.3363 | 0.5082 |
| CLTA | -0.6811 | -0.4791 | -0.8579 | 0.6141 |
| GGA1 | -0.9354 | -1.8144 | -0.1663 | 0.6159 |
| AP1G2 | -0.5424 | -0.3752 | -0.7096 | 0.7309 |
| CLTB | -1.2799 | -1.4830 | -1.0768 | 0.7726 |
| AP3B1 | -0.6350 | -0.2952 | -0.8898 | 1.0000 |
| GGA3 | -0.5631 | 0.1132 | -1.2394 | 1.0000 |
| AP1S1 | -0.4277 | -0.6314 | -0.2494 | 1.0000 |
| DNM1 | -0.3980 | -0.7538 | 0.0087 | 1.0000 |
| DNAJC6 | -0.3319 | -0.9135 | 0.2497 | 1.0000 |
| DNM3 | -0.3257 | -0.4909 | -0.2018 | 1.0000 |
| AP2S1 | -0.3014 | -1.1190 | 0.5162 | 1.0000 |
| CLTC | -0.2952 | -0.9897 | 0.3993 | 1.0000 |
| AP3D1 | -0.0420 | 0.0679 | -0.1382 | 1.0000 |
| SNAP91 | 0.2111 | -0.0651 | 0.4873 | 1.0000 |
| PI4K2A | 0.3820 | 0.3825 | 0.3815 | 1.0000 |
| VPS35 | 0.4090 | 0.5897 | 0.2025 | 1.0000 |
| ARFIP1 | 0.4159 | -0.2041 | 0.8809 | 1.0000 |
| ARF5 | 0.4356 | 1.1830 | -0.3119 | 1.0000 |
| AP3B2 | 0.4753 | 0.0492 | 0.8482 | 1.0000 |
| AP3M1 | 0.5321 | -0.3655 | 1.4296 | 1.0000 |
| AP1G1 | 0.6736 | 1.3136 | 0.0337 | 1.0000 |
| ARFIP2 | 0.7438 | 0.3163 | 1.1179 | 1.0000 |
| AP2B1 | 1.0545 | -0.0139 | 2.1229 | 1.0000 |

$P < 0.05$

c

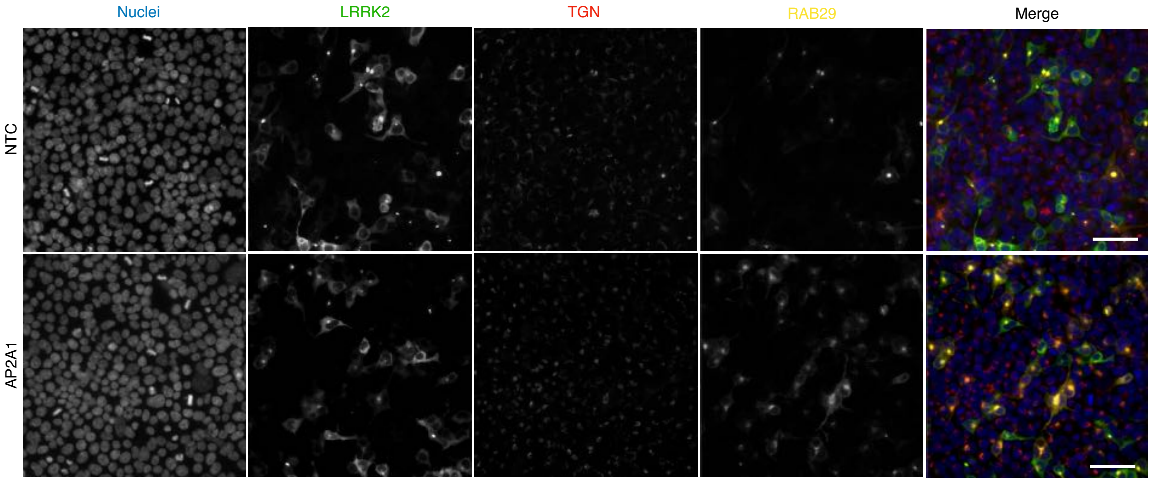

Supplementary table 3.

| Accession | MASCOT Score | Mass | Peptide matches | Significant peptide matches | Sequence matches | Significant sequence matches | emPAI | Protein | Gene | Tissue |
|---|---|---|---|---|---|---|---|---|---|---|
| AP2B1_MOUSE | 1955 | 104516 | 100 | 63 | 49 | 35 | 4.26 | AP-2 complex subunit beta | AP2B1 | Brain |
| AP2A2_MOUSE | 1215 | 103951 | 79 | 41 | 44 | 27 | 2.68 | AP-2 complex subunit alpha-2 | AP2A2 | Brain |
| AP1B1_MOUSE | 1022 | 103869 | 73 | 39 | 43 | 30 | 2.84 | AP-1 complex subunit beta-1 | AP1B1 | Brain |
| AP2M1_MOUSE | 487 | 49623 | 47 | 23 | 26 | 16 | 3.61 | AP-2 complex subunit mu | AP2M1 | Brain |
| DYN3_MOUSE | 393 | 97130 | 40 | 12 | 28 | 8 | 0.48 | Dynamin-3 | DNM3 | Brain |
| AP2A2_MOUSE | 321 | 103951 | 17 | 9 | 15 | 9 | 0.44 | AP-2 complex subunit alpha-2 | AP2A2 | Kidney |
| ARF4_MOUSE | 245 | 20384 | 15 | 8 | 8 | 6 | 3.18 | ADP-ribosylation factor 4 | ARF4 | Kidney |
| ARF5_MOUSE | 224 | 20517 | 19 | 11 | 9 | 6 | 3.15 | ADP-ribosylation factor 5 | ARF5 | Kidney |
| ARF1_MOUSE | 210 | 20684 | 13 | 10 | 9 | 7 | 3.11 | ADP-ribosylation factor 1 | ARF1 | Brain |
| ARF6_MOUSE | 172 | 20069 | 8 | 5 | 6 | 4 | 1.3 | ADP-ribosylation factor 6 | ARF6 | Kidney |
| AP3B2_MOUSE | 131 | 119118 | 20 | 4 | 19 | 4 | 0.15 | AP-3 complex subunit beta-2 | AP3B2 | Brain |
| ARF4_MOUSE | 120 | 20384 | 6 | 6 | 5 | 5 | 1.77 | ADP-ribosylation factor 4 | ARF4 | Brain |
| VPS35_MOUSE | 115 | 91655 | 6 | 2 | 5 | 1 | 0.05 | Vacuolar protein sorting-associated protein 35 | VPS35 | Brain |
| AP1S1_MOUSE | 115 | 18721 | 2 | 2 | 2 | 2 | 0.56 | AP-1 complex subunit sigma-1A | AP1S1 | Kidney |
| ARF6_MOUSE | 113 | 20069 | 5 | 3 | 4 | 3 | 0.86 | ADP-ribosylation factor 6 | ARF6 | Brain |
| CLH1_MOUSE | 104 | 191435 | 12 | 4 | 7 | 4 | 0.09 | Clathrin heavy chain 1 | CLTC | Kidney |

Supplementary Figure 4.

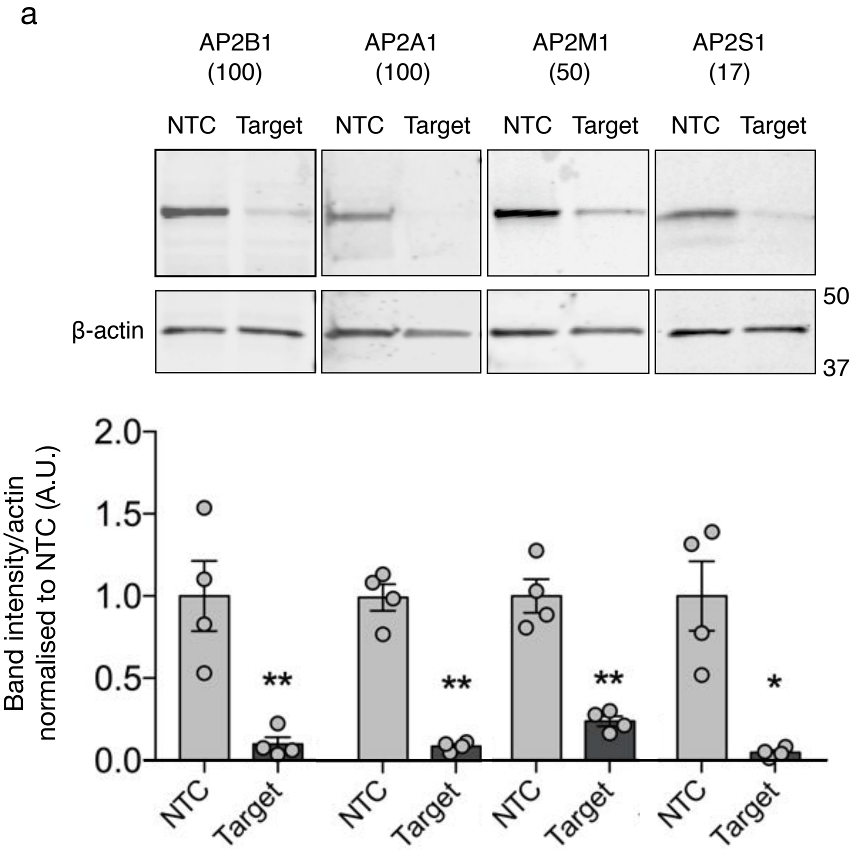

Supplementary Figure 5.

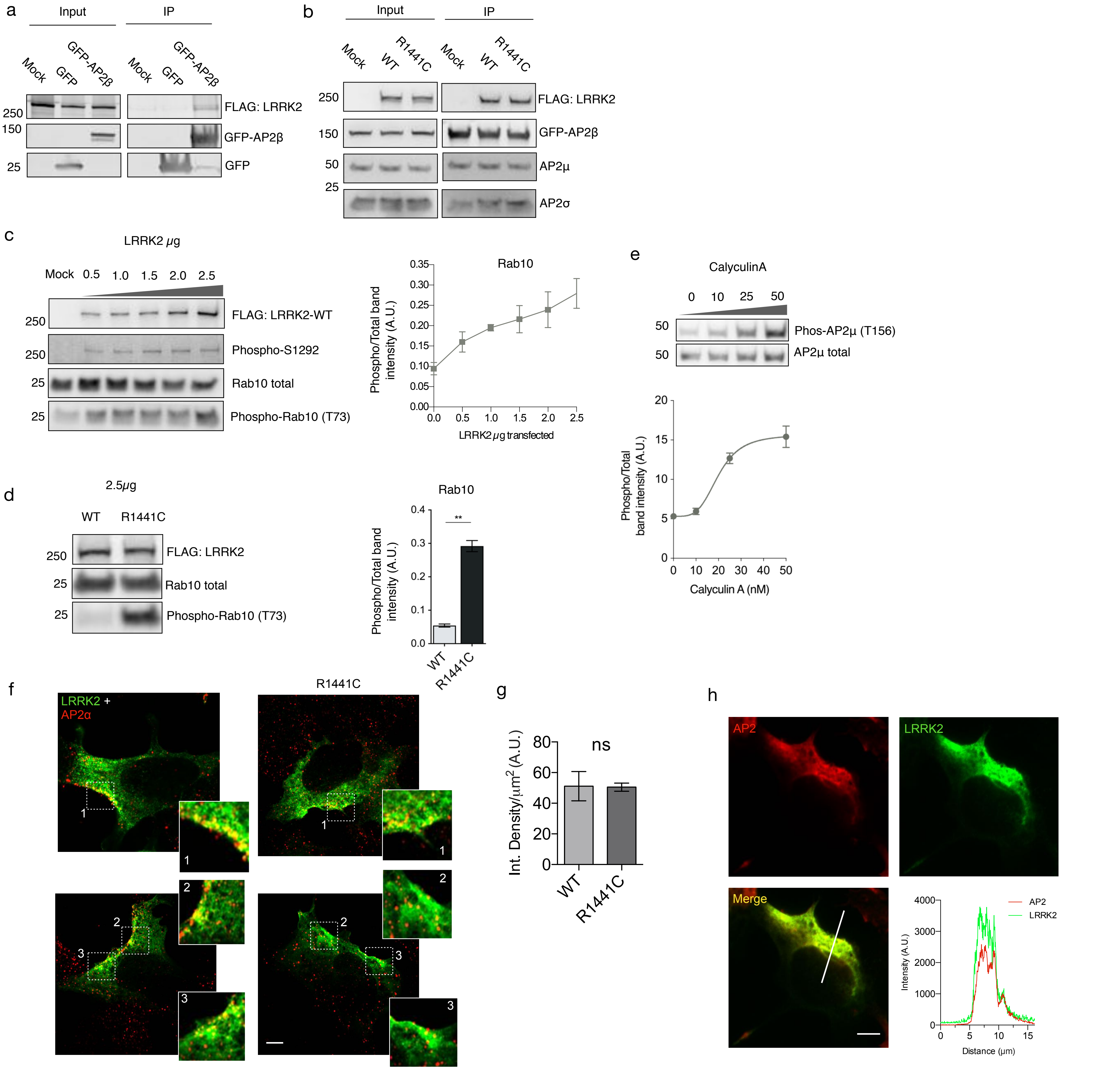

### Supplementary Figure 6.

a

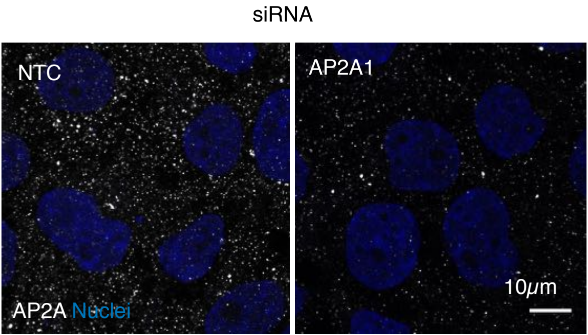

Supplementary figure 7.

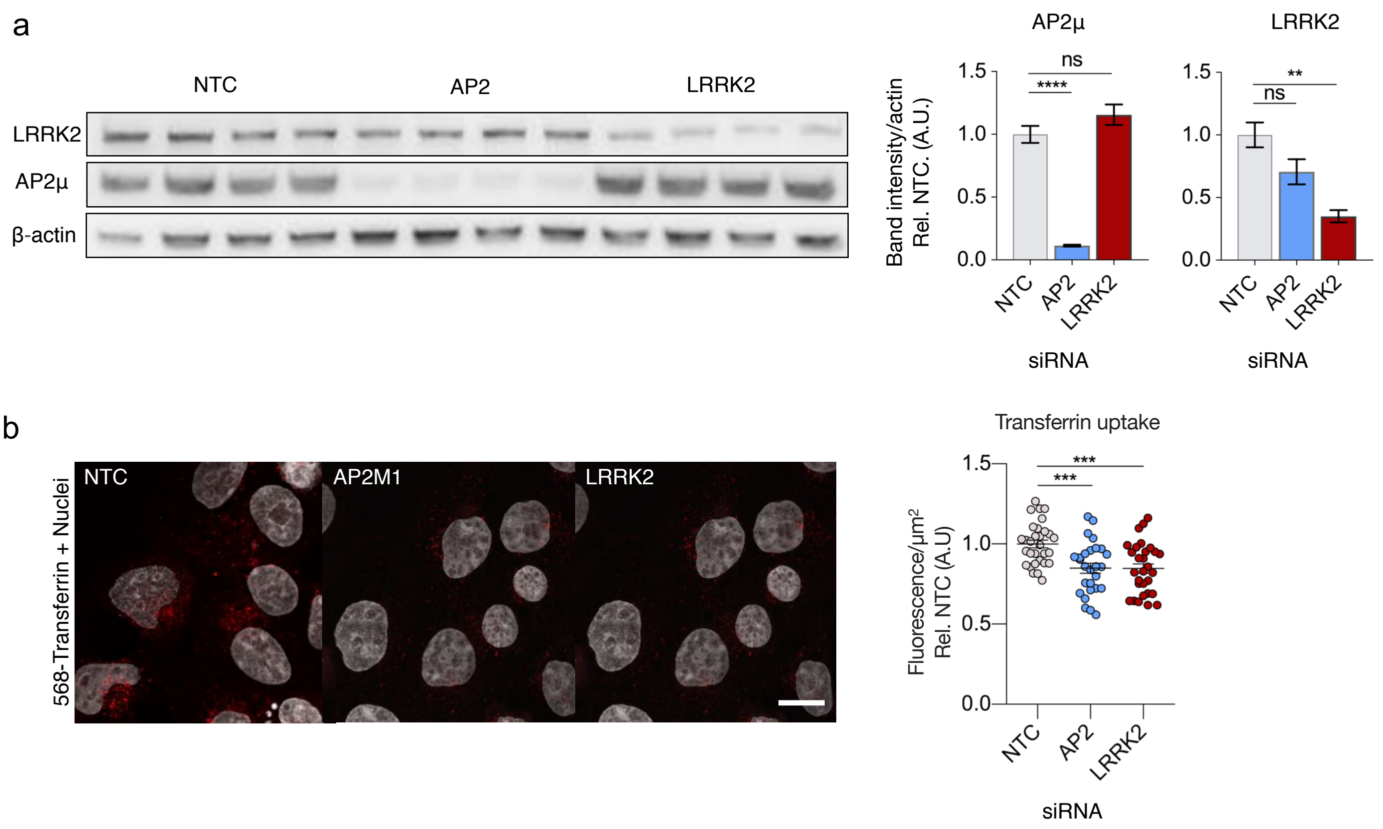

Supplementary figure 8.

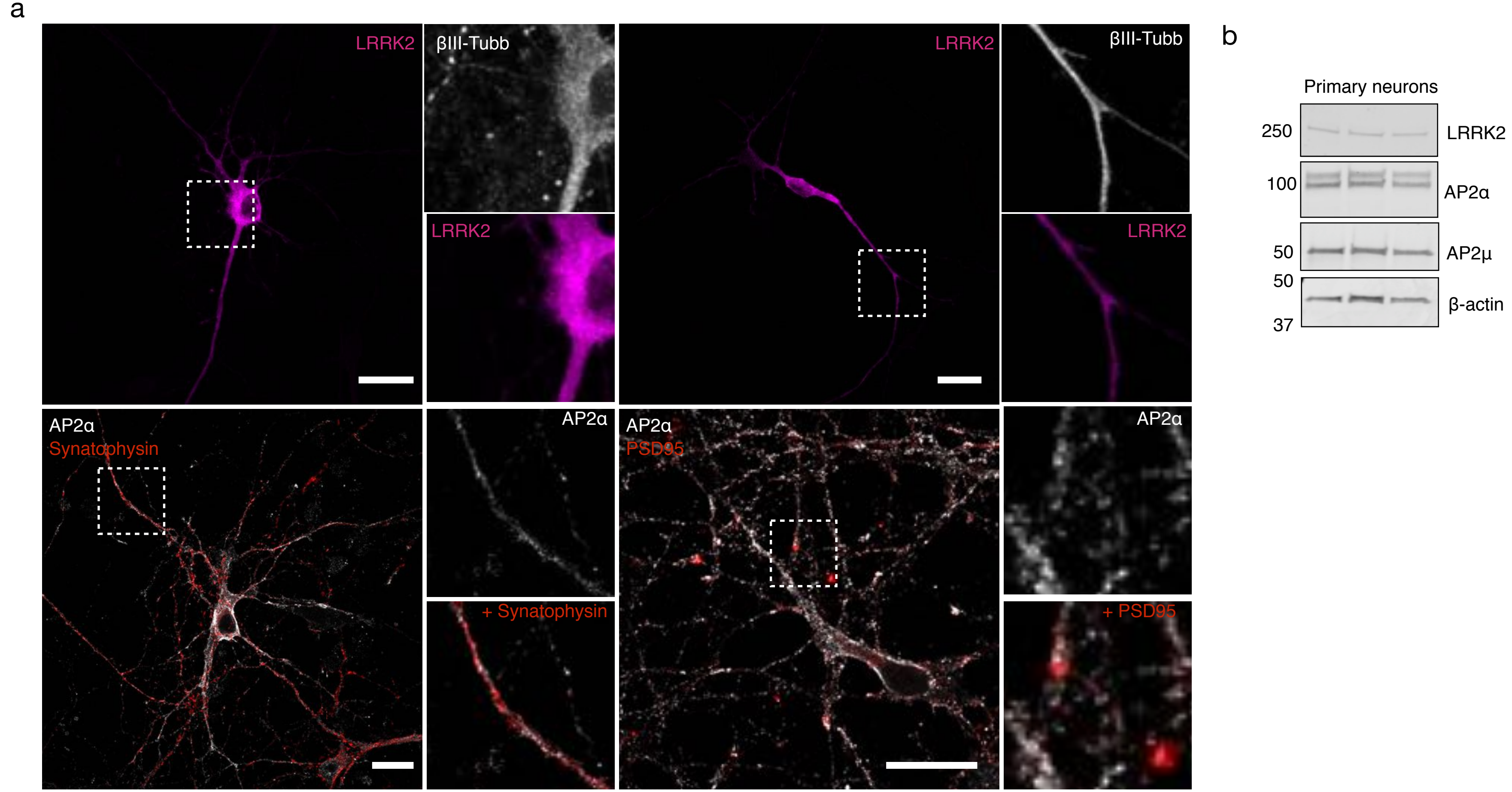

Supplementary table 9.

| Study | Cases (N) | Controls (N) | Total (N) | Case age at onset (mean , SD in years) | Control age at last exam (mean, SD in years) |
|---|---|---|---|---|---|
| Baylor UMary Parkinson’s | 789 | 195 | 984 | 64.9 (10.11) | 65.45 (8.31) |
| Finnish Parkinson's | 386 | 493 | 879 | 55.27 (5.64) | 92.35 (3.86) |
| McGill Parkinson's | 583 | 906 | 1489 | 65.71 (9.78) | 55.79 (10.69) |
| Oslo Parkinson's Disease Study | 476 | 462 | 938 | 50 (4) | 61.85 (11.06) |
| Spanish Parkinson’s | 1920 | 1164 | 3084 | 60.07 (12.70) | 69.02 (9.95) |
| Vance (dbGap phs000394) | 621 | 303 | 924 | NA | 81.88 (12.73) |
| Tubingen | 741 | 944 | 1685 | 55.76 (11.55) | 47.42 (12.38) |
| TOTAL | 5516 | 4467 | 9983 | | |
